# Equiluminant Border Ownership Cells as a Missing Link in Color Form Perception

**DOI:** 10.64898/2025.12.07.692838

**Authors:** Paria Mehrani, John K. Tsotsos

## Abstract

We propose the existence of a sub-population of border ownership neurons that signal figure side along equiluminant borders, those defined purely by color contrast. We postulate that these neurons, similar to primate equiluminant V1 [26] and V4 [5] cells, are strongly activated by equiluminant contrast on their preferred side, with diminished response for strictly luminance contrast. To test this hypothesis, we propose a hierarchical and mechanistic model that explains how equiluminant border ownership signals can be achieved in the ventral stream. Our simulation results suggest that model equiluminant neurons are highly color-selective while exhibiting similar orientation and border ownership selectivity as their luminance counterparts. Additionally, our model equiluminant cells indeed exhibited the above postulated pattern of responses. We suggest that these neurons are the missing links in the equiluminance channels in the ventral stream that transform equiluminant oriented edge signals in V1 to equiluminant object-centered shape representations in V4. Our findings point to the importance of re-examining the border ownership neuron population to further detail their role in equiluminant channels in the ventral stream.

## 1 Introduction

Contrast in luminance and color creates edges in an image that signal discontinuities in depth, reflectance, surface orientation and illumination [30], information conveying properties of the observed scene and the objects in it. For example, a subset of depth discontinuity edges correspond to borders of closer objects occluding either the ground or other objects, giving rise to contours that lead to object shape. Accordingly, edge information forms the foundation of shape representation and form perception in both biological [24] and machine vision [30]. In primates, edge signals are processed through a hierarchy of cortical areas, where they are progressively transformed into more complex shape representations that support object perception. In V2, the integration of local edge signals gives rise to higher-order features such as junctions and corners. Further along the ventral stream, neurons in V4 exhibit selectivity for object-centered shape components, including convex and concave parts [38]. Such object-centered representations depend critically on figure–ground assignment, *ie*., distinguishing the inside from the outside of an object along the occlusion borders. This computation is supported by border ownership (BO) neurons [59], which explicitly encode the side of an occlusion border that belongs to the object. Thus, BO cells are considered as a key source of inside–outside information to downstream shape-selective neurons in V4 [34, 39, 53, 35].

While most perception research has been based on the view that the form subsystem in the primate visual cortex is achromatic and that neurons selective to oriented luminance edges are the major contributors to form perception (See [17, 18, 49, 47] for reviews), Hansen and Gegenfurtner [20] demonstrated that equiluminant edges defined purely by color contrast are statistically independent of and not rarer than luminance edges in natural scenes. For example, the natural scene image shown in Figure 1 includes objects whose shapes are defined solely by equiluminant edges. They further suggested that equiluminant and luminance edges jointly define figure borders, supporting a coherent perception of object shape.

**Figure 1:**
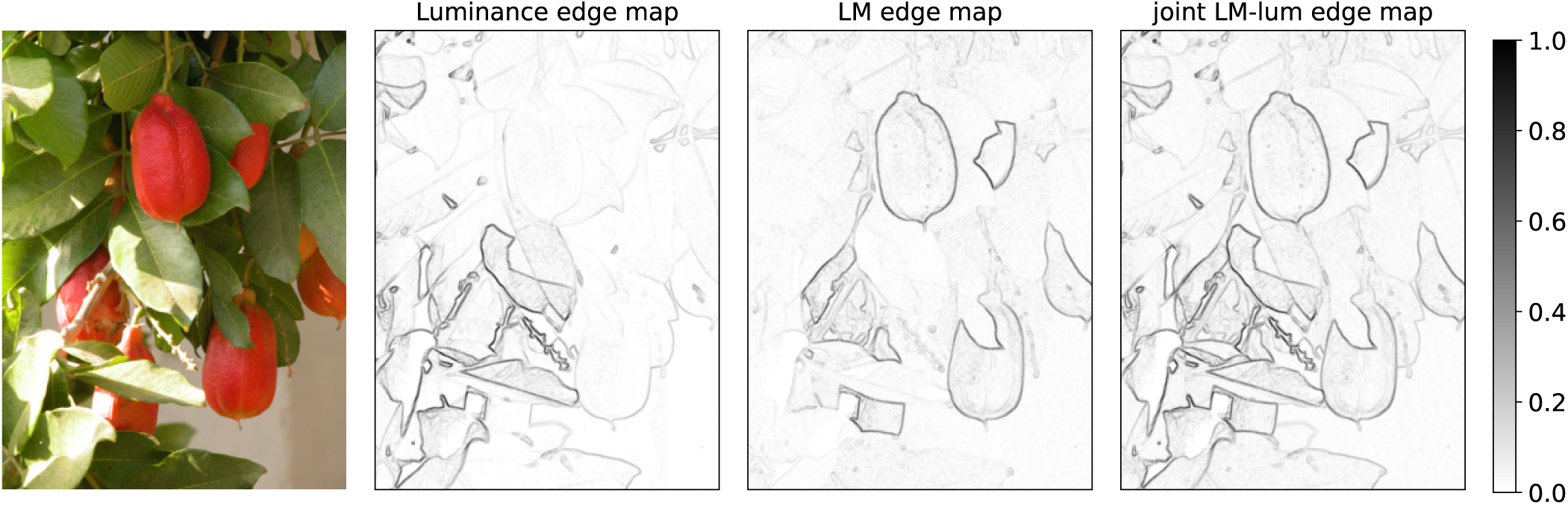
A natural scene example from [36] demonstrating existence and presence of luminance and equiluminant edges (re-produced from [20]). We followed the same procedure as defined in [20] to convert RGB images to LMS cone activation, computing luminance maps and extracting edges using 3×3 Sobel operators. From left to right, original image, luminance edge map, LM edge map, joint LM and luminance edge maps. Comparing luminance and LM edge maps highlight a disjoint set of edges between these maps. This example demonstrates that equiluminant edges, independently defined from luminance ones, exist in natural scene, highlighting the importance of accounting for these features in form perception. The joint map on the right provides a more complete representation of the input edges than the individual luminance and LM maps.

Evidence on edge- and shape-selective color cells in primate V1 and V4 supports the presence of an equiluminant subsystem contributing to object shape perception. In particular, several studies suggested an integrated, not a segregated, color-form processing system, reporting a spectrum of color and form selectivities [26, 25, 27, 16, 46, 5, 6, 7] (See [48, 49, 44] for reviews). A sub-population of neurons in this spectrum represent shapes defined solely by equiluminant edges. Specifically, a distinct sub-population of V1 neurons (about 12%) strongly respond to equiluminant oriented borders and weakly to luminance ones [26, 16]. These equiluminant orientation-selective cells exhibit similar spatial processing properties as those of luminance cells, making them early stage candidates for color boundary detection and consequently color object perception [48]. Similarly in V4, equiluminant shape-selective cells were found by Bushnell *et al*. [5, 6] who reported that these neurons responded strongly to shapes defined by equiluminant contrast and weakly to those defined by luminance ones. These cells, comprising about 22% of the studied population, showed similar object-centered shape selectivities as luminance shape cells in V4 [37], depending on inside-outside information along equiluminant occlusion borders. Although color selectivity was found in about 40% of BO neurons [59, 15], BO responses to equiluminant stimuli are unknown, leaving the cortical representations of figures defined by equiluminant edges an open question. Here, we propose such cells must exist to complete the representation of figure-ground assignment in the ventral stream and introduce a hierarchical computational model of equiluminant border ownership (EBO) cells that explains how these representations can be achieved in the ventral stream. Additionally, we predict their cortical characteristics, *ie*., strong responses to their preferred equiluminant stimuli and weak activations in the presence of luminance input.

Our proposed model builds on prior findings suggesting similar mechanisms in equiluminant and luminance form processing pathways [7] and extends the Recurrent Border Ownership (RBO) network [33] that performs BO assignment across luminance borders. The RBO is a hierarchical computational network modeling neurons in both ventral and dorsal streams. The model ventral stream implements simple, complex and BO cells in V1 while the model dorsal stream is comprised of dorsal simple V1 cells and MT neurons. Previous studies have shown that BO cell responses depend not only on local features within their classical receptive fields but also on contextual information [59, 58, 51]. In RBO, model MT neurons with their large receptive fields provide this contextual information, imparting an initial figure-side preference to model BO cells. The BO signal is further enhanced through lateral interactions among neighboring model BO neurons (See Supp. Information and [33] for further details and the choice of architecture).

Our proposed extension, whose architecture is depicted in Figure 2 and we call eRBO (equiluminant RBO), follows the same underlying mechanisms and architecture as RBO. Input to eRBO consists of Long (L), Medium (M) and Short (S) cone activations. To model luminance pathways in primate visual cortex, L and M cone activations are linearly combined into a luminance map [20] passed to luminance pathways in eRBO. The computations in luminance pathways in eRBO are identical to those in the original RBO network. Model BO cells in RBO inherit their local feature selectivities from model simple and complex cells. To model equiluminant edge selectivity in the extended network’s ventral cells, we implemented orientation-selective equiluminant simple cells according to the findings of Johnson *et al*. [26] who reported equiluminant double-opponent cells in V1. Determined by cone contributions to our model equiluminant V1 cells, four types of color selectivities were modeled in eRBO. Responses of model equiluminant simple neurons are combined in equiluminant complex cells that send a feed-forward signal to EBO cells in our model ventral stream. Importantly, through our suggested mechanisms, we propose that the onset of equiluminant oriented edge responses in primate EBO cells is initiated by feed-forward signals from equiluminant orientation-selective V1 neurons [26], linking equiluminant edge signals in V1 to more complex equiluminant shape selectivities observed in V4.

**Figure 2:**
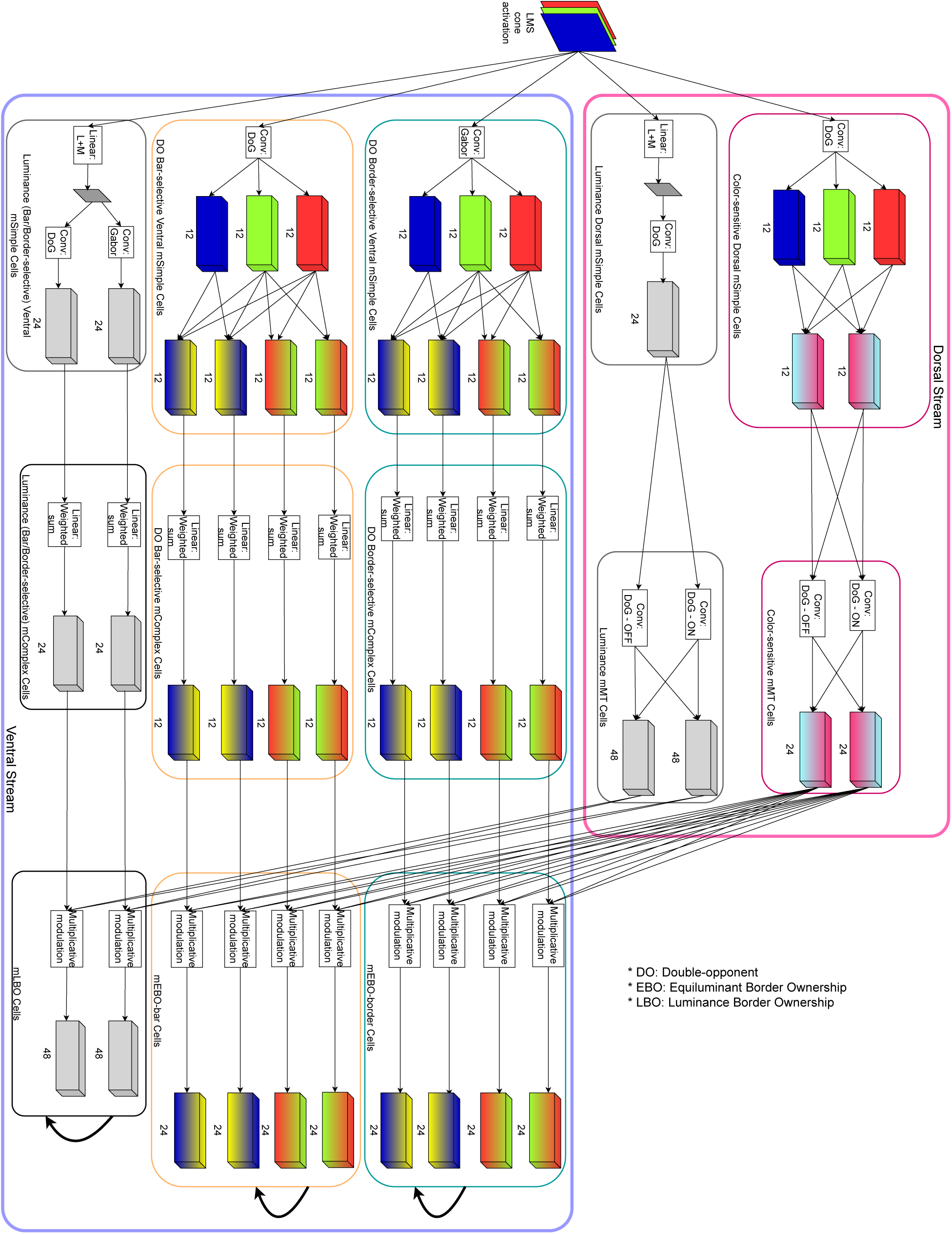
The eRBO model architecture consisting of a two-stream network modeling neurons in ventral and dorsal pathways. The letter “m” preceding neuron types in the figure and caption denotes model cells. Input to the model are L, M, and S cone activations. In the luminance sub-populations in both streams, the luminance map is computed as a linear combination of L and M cone activations. The luminance sub-populations follow the same computations as those described in [33]. The color sub-populations in eRBO were modeled based on color-sensitive neuron data in the primate dorsal stream [19, 9, 45, 4, 8, 9]. The equiluminance signals in the ventral stream start in equiluminant ventral model simple cells that were modeled according to double-opponent cells in primate V1 following [26]. Responses of these model neurons are aggregated to generate equiluminance model complex cell responses that send a feed-forward signal to mEBO cells in the ventral stream. Color-sensitive dorsal mMT neurons multiplicatively modulate mEBO cells and provide global context from outside their classical receptive fields to these cells. Responses of mEBO cells, similar to their luminance counterparts, are then enhanced with lateral modulations within local neighborhoods. This latter step is conducted among individual sub-populations - for example, mEBO-border.

In RBO, the early emergence of asymmetric responses in model BO cells was driven by recurrent input from on- and off-center model MT cells, whose receptive fields featured elongated excitatory and inhibitory regions [12]. The summed responses of these neurons across all orientations and from multiple surround distances on the preferred side signaled the presence of closed shapes on either side of an edge, hence providing figure context to model BO cells. Here, we modeled dorsal cell responses following [19, 9, 45, 4, 8, 56], studies that challenged lack of color responses in the dorsal pathway and reported color sensitivity in this stream. Accordingly, we modeled color-sensitive dorsal simple V1 and MT cells with the same spatial processing properties as those of luminance dorsal neurons in the RBO but set their color response properties based on [4]. In eRBO, dorsal modulation is applied to model BO neurons with similar color/luminance selectivity followed by relaxation labeling performing lateral interactions among luminance and equiluminant sub-populations separately. As a result, contextual color figure information conveyed by color-sensitive dorsal cells on the preferred side of a model EBO cell initiates the BO signal along equiluminant edges, complementing the BO signal carried by LBO cells along luminance-induced figure borders. In RBO, two types of local feature selectivities were modeled: selectivity to borders between light-dark regions and selectivity to light/dark bars between two dark/light regions. Here, we modeled the same two types of local feature selectivity and extended the pattern to our model double-opponent simple cells (See Supp. Information, Figure 2 for details). In eRBO, all neuron types with bar and border selectivity are modeled at 4 spatial scales, 2 contrast polarities and 12 orientations.

Our simulation results confirm color response properties in model BO neurons as reported in [59, 15]: we report strong responses to color stimuli in our model EBO cells. Our results affirm comparable orientation and BO signal properties in both model EBO and luminance BO cells. However, advancing beyond the findings of [59, 15], we report that model EBO cells exhibit similar response patterns as those of primate equiluminant V1 and V4 cells [26, 5]. That is, our model EBO neurons respond strongly to shapes defined by equiluminant borders in the stimulus, and weakly to luminance defined borders. In contrast, model LBO neurons weakly responded to equiluminant stimuli, yet displayed pronounced activations to luminance stimuli. These findings highlight the distinct role each sub-population plays in forming a coherent shape representation in the visual cortex and the need for EBO representations to encode inside-outside information along equiluminant occlusion boundaries to V4 and higher visual areas. Despite extensive research on BO neurons [58, 59, 15, 41, 40, 54], to the best of our knowledge, this work is the first attempt to propose the existence of EBO cells, predicting their cortical characteristics, and providing a mechanistic account on the emergence of EBO signals in the primate visual cortex.

In what follows, we will make a distinction between our model and brain cells by a preceding “m” for model neurons, for example, “mV1” for model V1 cells.

## 2 Results

To examine the effects of border ownership in our model neurons, we created a stimulus set following [59], with example images shown in Figure 3a. To isolate luminance versus color influences on model neuron responses, we generated pure luminance images with luminance values set to those utilized in [59, 15]. We created color images with 44 sampled hues from the DKL space whose chromatic axes represent *L* − *M* and *S* − (*L* + *M* ) cone interactions [11]. We sampled hues at elevation=0 and contrast=1, with sampling separation determined by the just-noticeable-difference extracted from Witzel and Gegenfurtner [55] (their Figure 5). The sampled colors were assigned to one region in the image (figure or ground) and the other color was set to black to ensure same activation level (zero in this case) in all three cone types.

**Figure 3:**
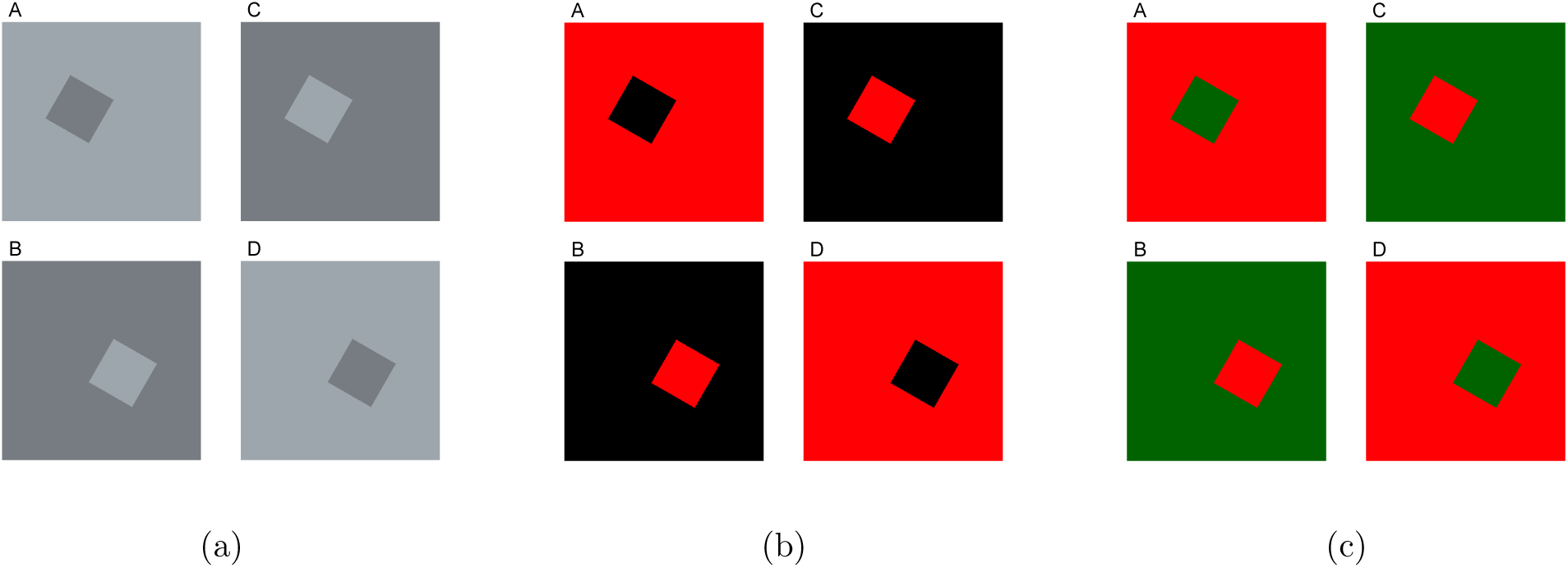
Border ownership stimuli example. 3a represents stimuli in gray, 3b demonstrate the color BO stimuli, and 3c shows the equiluminant stimuli with figures defined purely by color contrast between red and green regions. In all sets, the four images show a square figure with one side centered in the middle of each image. The images in each row (A,C and B,D) share the same figure side at opposite polarity contrast. Images in each column (A, B and C, D) share the same polarity (luminance or color) but have opposite figure direction. BO cell receptive fields fall on the figure side shared between the four images.

The results below summarize our investigation of the properties of mEBO cells. First, we show that aligned with the findings of equiluminant edge and shape selective neurons in primate V1 and V4 [26, 5], our mEBO neuron responses generated by the proposed mechanisms exhibit orientation and BO selectivities similar to those of mLBO cells. To assess color selectivity in mEBO neurons, consistent with the findings of [59, 15], we examined mEBO responses to color BO stimuli described above. Our primary hypothesis, however, is beyond the reported color selectivity in BO cells [59, 15] and concerns strong activation of mEBO cells in response to their preferred equiluminant stimuli and weak responses to luminance ones. To test this response pattern, and inspired by the red–green stimuli used to probe equiluminant double-opponent V1 cells [26], we created a set of equiluminant red–green BO stimuli, as illustrated in Figure 3c. In this stimulus set, figure borders are defined purely by color contrast, providing a controlled means of evaluating model EBO responses to equiluminant figures. Finally, we demonstrate that mEBO and mLBO cells together provide complementary and comprehensive figure-side information along both luminance and equiluminant defined borders of fruits and leaves in the natural scene shown in Figure 1.

### 2.1 Similar orientation and BO selectivities in mLBO and mEBO cells

We evaluated orientation and BO selectivity in mEBO and mLBO neurons to establish whether the two populations have identical shape processing characteristics, replicating a similar observation reported in equiluminant V1 and V4 cells [26, 5]. Specifically, if the two sub-populations share similar shape-processing properties, both encode shape along color or luminance contrast edges according to their local pattern preference. Orientation selectivity tests were conducted by identifying preferred color for each model neuron and recording responses to square sides oriented at 12 equally spaced orientations across 180^◦^. We measured orientation tuning bandwidth (half-width at half-peak following [26]) and orientation modulation index calculated as 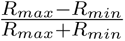 [15]. Additionally, BO selectivity was assessed using a BO modulation index defined as the normalized difference of responses to squares with preferred color and orientation presented on opposite sides of the border (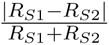 following [15], where *S*1*, S*2 represent figures on two sides of the border).

Orientation tuning bandwidth in our model luminance and equiluminant complex cells as well as mEBO and mLBO neurons was set to 15^◦^, indicating high levels of orientation selectivity in these cells. Similar orientation tuning bandwidth observed across model complex and mBO cells suggests that, as expected, both mEBO and mLBO neurons preserve the orientation selectivity that is transmitted to them through model complex cells.

Figure 4 illustrates orientation modulation index plotted against border modulation index for mLBO and mEBO cells. The two side histograms in this figure show a substantial overlap in both orientation and BO modulation index distributions across all model cells, confirming comparable orientation and BO selectivity properties in these cells. Both mLBO and mEBO cells are highly orientation-selective, with orientation modulation index values greater than 0.86. Additionally, the skewness in BO modulation index distributions for all model neurons toward values greater than 0.2, corresponding to responses twice as strong to preferred BO stimuli than non-preferred stimuli, suggests a distinctive figure-side preference across mEBO and mLBO cells.

**Figure 4:**
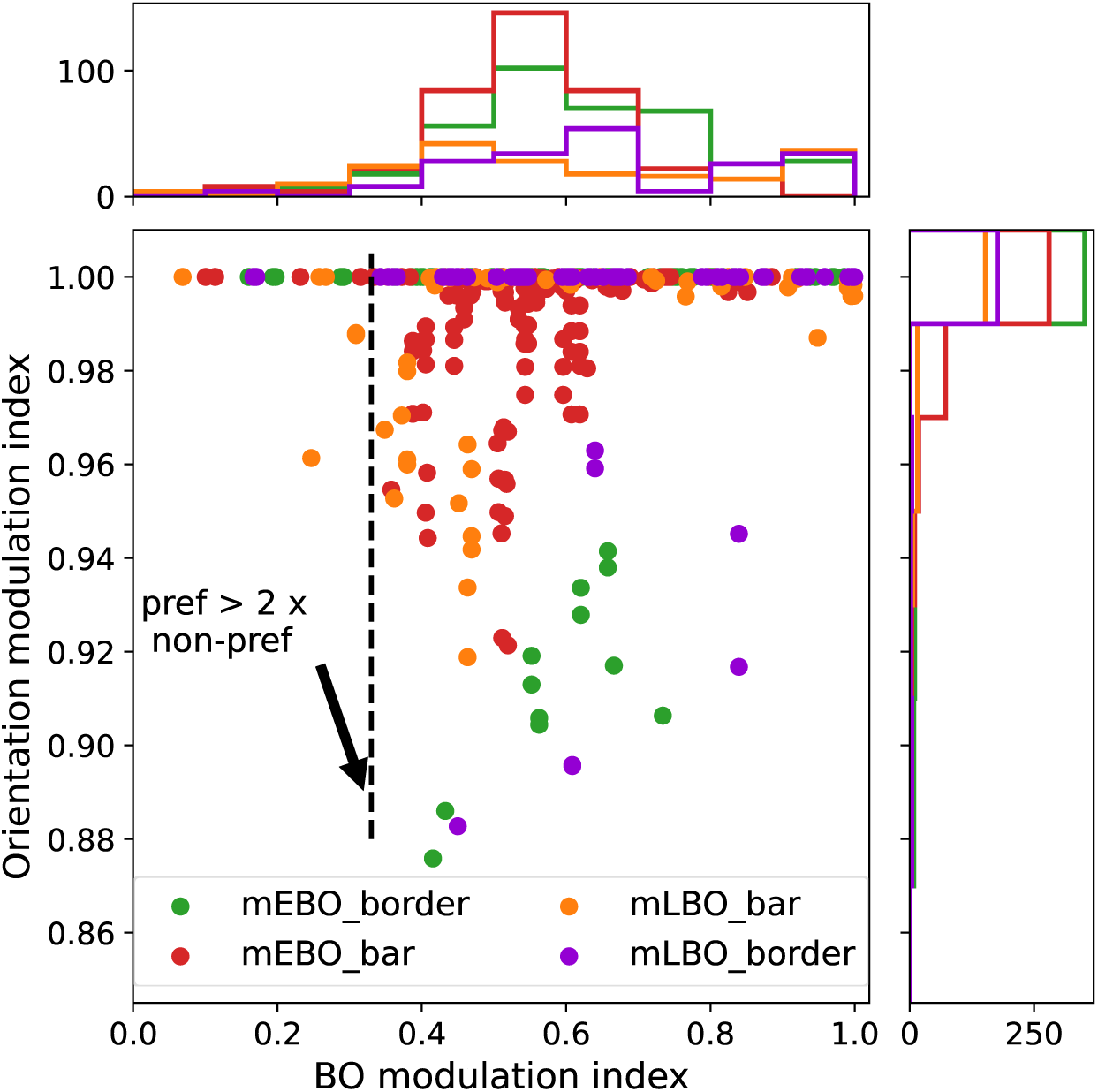
Orientation and BO selectivity analysis. The scatter plot represents orientation modulation index vs. BO selectivity in mEBO and mLBO neurons. Each cell type is represented by a unique color dot. The vertical dashed line demonstrates the BO selectivity index threshold above which responses to preferred stimuli are at least twice as strong as those of non-preferred stimuli. The side histograms above and on the right demonstrate the distribution of BO selectivity index and orientation modulation index respectively. These histograms indicate a substantial overlap in the distributions of the different neuron types, suggesting similar orientation and BO selectivities among mLBO and mEBO sub-populations.

### 2.2 High color selectivity in mEBO, not mLBO, neurons

We tested color selectivity in mEBO and mLBO neurons by measuring three quantities: color tuning bandwidth [28], color selectivity index and achromatic index [15]. While color tuning bandwidth, measured as half-width at half-peak, denotes the range of colors to which a neuron exhibits fairly strong responses, it does not express the strength of response to preferred color versus other color and luminance stimuli. To quantify the relative strength of responses to a preferred color, we computed a color selectivity index calculated as 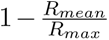, where *R_mean_, R_max_* represent mean and max responses to all color and luminance stimuli. Additionally, we computed an achromatic index as 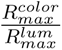, where 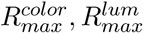 denote maximum response of a neuron to the most preferred color and luminance stimuli respectively. Smaller color tuning bandwidths together with greater color selectivity index and achromatic index values suggest high color selectivities in a neuron.

Figure 5a illustrates the color tuning bandwidth distributions for model equiluminant and luminance complex and mBO cells. The shift to smaller tuning bandwidths in equiluminant compared to luminance model complex and mBO neurons is evident. Specifically, mean color tuning bandwidth is at 74^◦^ and 64^◦^, near orthogonal hues, indicating relatively strong responses to a wide range of hues in model luminance complex and mLBO neurons. In contrast, model equiluminance complex and mEBO cells demonstrated narrower tunings with mean bandwidth at 42^◦^ and 34^◦^ respectively, suggesting higher levels of color selectivity in these cells.

**Figure 5:**
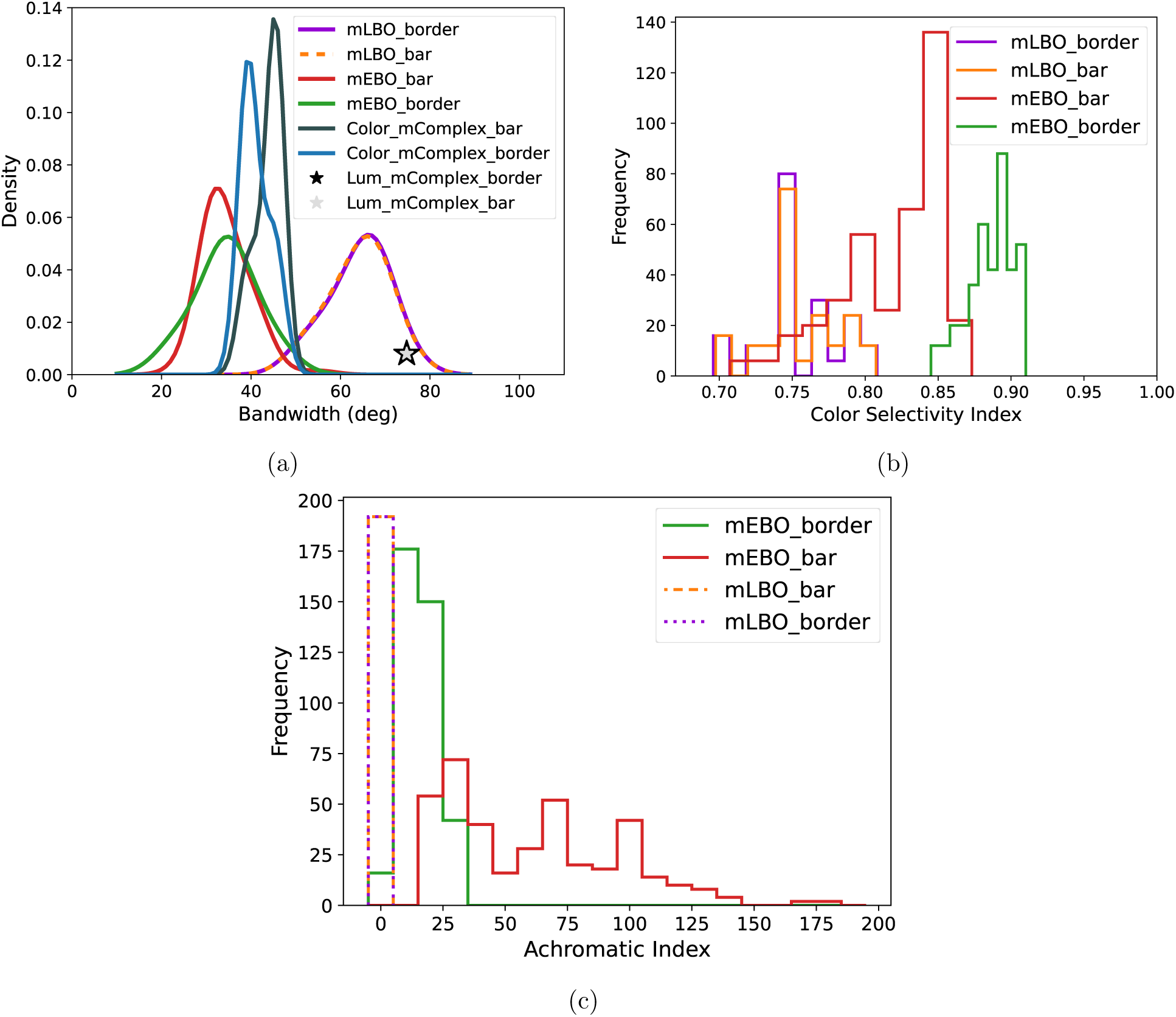
Color selectivity analysis. 5a Color tuning bandwidth in complex and BO sub-populations in the model. Note that equiluminant cells, both mComplex and mEBO, are highly color selective compared to luminance mComplex and mLBO cells. 5b Color Selectivity Index plotted for mLBO and mEBO sub-populations demonstrates the shift to larger color selectivity index value in mEBO neurons compared to mLBO cells, suggesting color selectivity in the mEBO sub-population. 5c The achromatic index histograms for both mEBO and mLBO neurons. The dashed bar edges indicate locations where bar edges of different neuron types overlap. While color selectivity index indicates color selectivity, achromatic index teases apart stronger responses to color stimuli vs luminance by computing the ratio of the response to the most effective chromatic stimuli over the response to the most effective achromatic stimuli. The histogram shows that the mLBO sub-population bars center at 0, indicating weak responses to color compared to luminance stimuli. In contrast, the significant shift in mEBO histograms suggests markedly stronger responses, in some cases by a factor of up to 175, to color than luminance stimuli. Due to the substantial overlap among the bars in Figure 5c, some bars are drawn with dashed or dotted edges to make the overlap visible.

The color selectivity index histograms in Figure 5b show greater values in mEBO than mLBO cells indicating a smaller ratio of 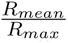 and therefore, higher selectivity in mEBO cells. However, the color selectivity index mixes responses to all colors (preferred and non-preferred) and luminance stimuli in computing *R_mean_*. To better tease apart the relative strength of neuron responses to color versus luminance stimuli, we measured achromatic index in mEBO and mLBO neurons. The achromatic index histograms plotted separately for the various mBO cell types in Figure 5c clearly show values near 1 for mLBO cells, indicating similar response strength to their most preferred chromatic and achromatic stimuli. In contrast, achromatic index values larger than 1, reaching up to ≈175, in mEBO cells reflect their strong selectivity to chromatic stimuli. Note, however, that our results do not indicate coding for hues in mEBO neurons but a pronounced preference for color stimuli. In fact, the measured color tuning bandwidth in mEBO cells is larger than tuning bandwidths reported in hue-selective cells [32, 28]. Finally, the three measurements together highlight lack of color selectivity in mLBO cells.

### 2.3 mEBO cells respond strongly to equiluminant, not luminance, stimuli

Our simulation results to this point indicate stronger responses to color stimuli in mEBO than mLBO neurons. Yet their responses to equiluminant and pure luminance stimuli remain unexamined. To address if mEBO cells exhibit similar responses as equiluminant shape cells in primate V1 and V4, we designed two sets of border ownership stimuli with equiluminant red-green (with elevation=0, contrast=1 and hues at 0 and 180 for red and green) and pure luminance regions, following [26]. Examples of the equiluminant and luminance stimuli are shown in Figure 3c and 3a respectively. We recorded mEBO and mLBO responses and normalized each neuron’s activation by its maximum response across equiluminant and luminance stimuli. Then, we measured the difference of responses to figures on opposite sides of the border as a quantitative measure of the border ownership signal for the neuron between the two stimulus types. The response differences in mEBO and mLBO neurons depicted in Figure 6a are visibly distinct: mLBO cell responses are populated along a vertical line in the two left quadrants of the scatter plot and mEBO cells are mostly along the horizontal line in the two bottom quadrants. Specifically, whereas mLBO cells respond similarly to opposite figure sides in equiluminant red-green stimuli, they exhibit relatively large response differences when presented with luminance border ownership stimuli. In contrast, mEBO response differences to luminance stimuli are near zero for most cells but span a wide range of values with equiluminant RG stimuli. This pattern in responses mirrors those of equiluminant V1 and V4 neurons in primates [26, 16, 5, 6].

**Figure 6:**
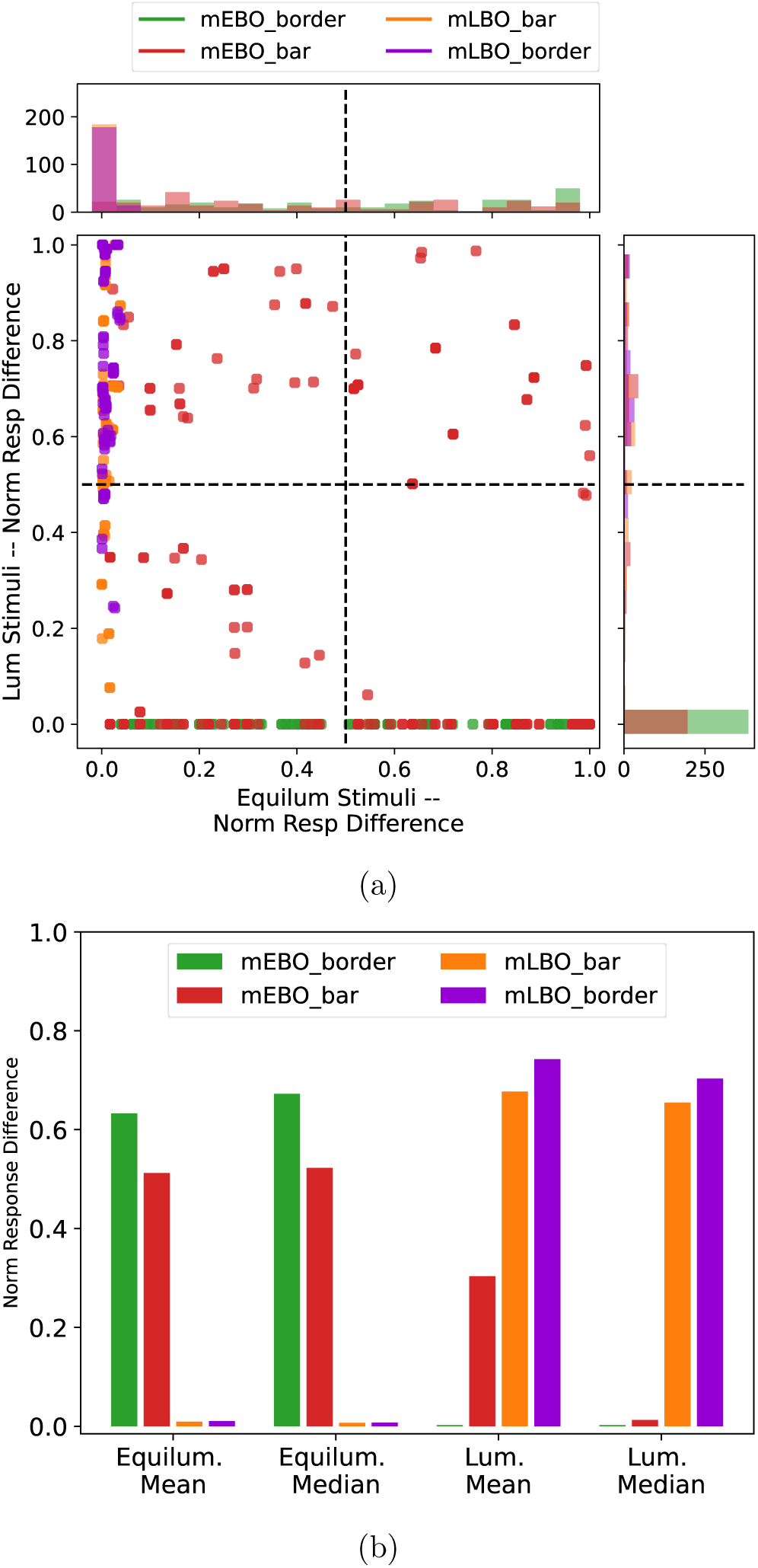
Equiluminance response analysis. 6a We computed mEBO and mLBO responses to luminance and equiluminant stimuli, normalized by maximum activation across stimuli. The difference of responses shown in the scatter plot indicates border ownership signal strength. Each dot in the figure represents a neuron type with similar colors employed in the two histograms above and on the right of the scatter plot. The mLBO population responses formed a vertical line on the left side of the scatter plot, indicating activation in presence of luminance stimuli while remaining unresponsive when figure is defined by color contrast. The two histograms confirm this pattern in mLBO responses. Note that bars in these histograms are plotted with 50% opacity to show overlapping distributions. For example, in the histogram on the right, the red and green bars overlap at 0. In contrast, mEBO neurons signal border ownership in presence of equiluminant stimuli and are mainly blind to luminance ones. Although some mEBO cells demonstrated relatively strong responses to luminance stimuli (≥ 0.5), those are at the 65th percentile. The histograms of mEBO neurons also confirm that the majority of these cells signal border ownership along equiluminant borders. 6b The mean and median of responses plotted separately for mEBO and mLBO populations to luminance and equiluminance stimuli. Note that mean and median of populations confirm the opposite pattern of responses among these two sub-population of model BO cells.

In Figure 6a, some mEBO cells exhibit relatively large response differences (≥ 0.5) to luminance stimuli (The red dots above the horizontal dashed line in the scatter plot). This population of mEBO neurons (all with S vs. LM selectivity) is at the 65th percentile of response differences. Therefore, the majority of mEBO cells exhibit weak border ownership signals to pure luminance stimuli. The mean and median of response difference distributions, presented in Figure 6b, confirm the observation that mEBO cells, similar to primate equiluminant V1 and V4 neurons, strongly respond to equiluminant stimuli but have weak to no activation when presented with pure luminance one. In contrast, mLBO cells are blind to equiluminant stimuli and only respond to figures defined by luminance borders.

### 2.4 mEBO and mLBO cells collectively represent figure ownership in natural scenes

The two neuronal populations of mEBO and mLBO cells exhibit opposite response patterns to equiluminant and luminance stimuli. With independently defined luminance and equiluminant figure borders in natural scenes [20], prevalence of both neuron type in vision systems for reliable shape representations becomes essential. In other words, lack of either neuronal type in a vision system would render an incomplete picture of shape signals in the observed scene, leading to unreliable shape representations in higher visual areas and impaired figure-ground organization.

Figure 7 demonstrates the necessity of integrating both representation type for effective figure-ground organization and equiluminant shape representations [5] in natural scenes. Following the example in Figure 1, we illustrated mEBO and mLBO responses to the input natural image separately across the four BO scales implemented in our model. Specifically, each pixel in these maps indicates the activation of the winning mLBO or mEBO cell whose receptive field center coincides with the pixel. In Figure 7, the BO cell with the highest activation in the sub-population was used as the winning BO cell for plotting. The ownership direction is color-coded, specified with the color-wheel arrows below the input image in Figure 7. The strength of response is indicated by saturation of the color with darker colors corresponding to stronger activations. Note that neuron activations in mLBO and mEBO maps coincide with the edges in luminance and LM maps in Figure 1. This outcome is expected as mBO neurons receive feed-forward signals from model luminance and equiluminant orientation-selective cells. However, the mLBO and mEBO maps in Figure 7 contain additional information: ownership direction obtained through contextual information from outside their classical receptive field, relayed by luminance or color-sensitive mMT neurons, respectively, followed by lateral interactions. Additional examples can be found in Supp. Information.

**Figure 7:**
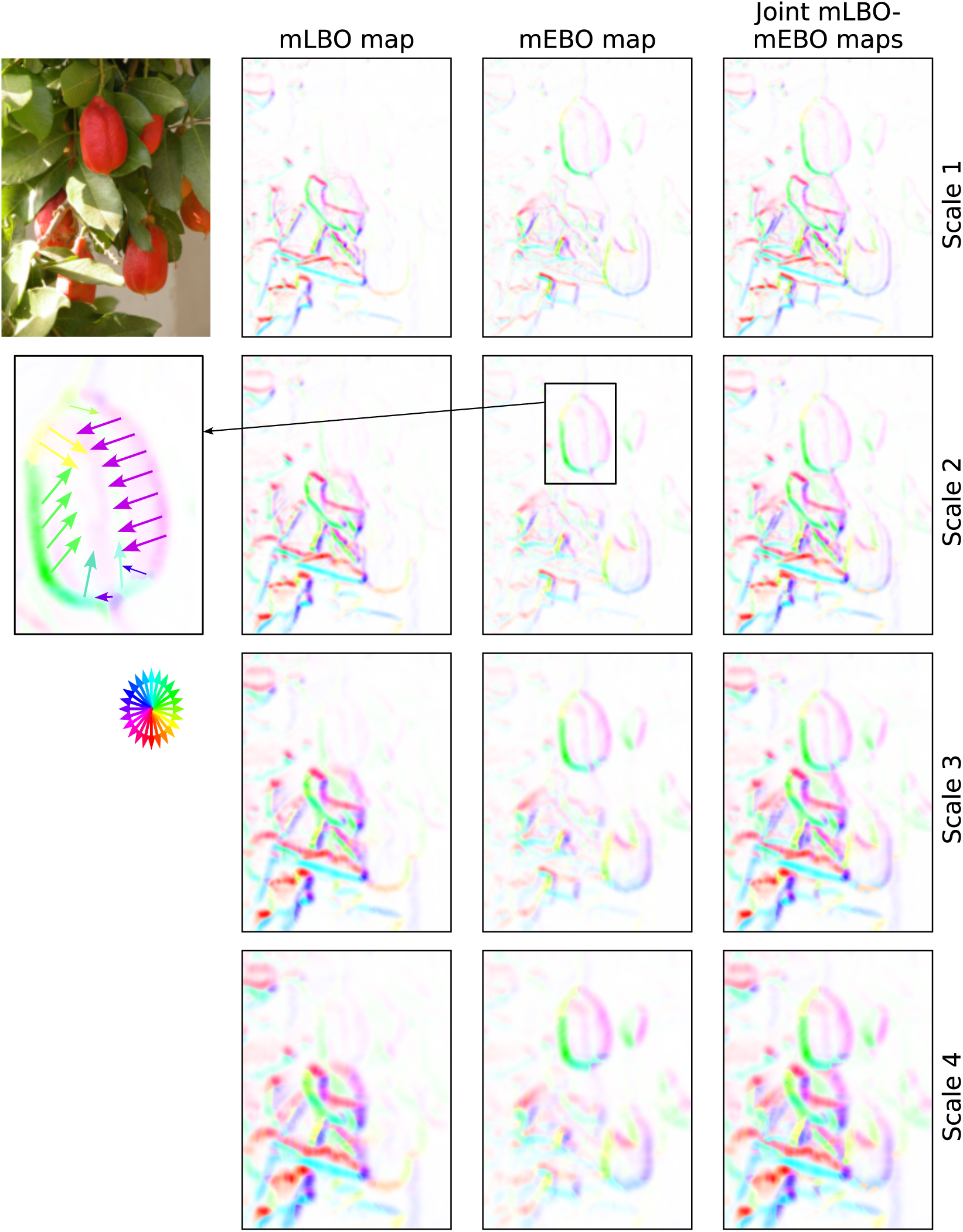
Coding of BO in a natural scene example. We plotted the LBO and EBO maps when the natural scene example of Figure 1 was given as input to eRBO. Each pixel in these maps represents response of the winning BO neuron whose receptive field centered on the pixel. The BO cell with strongest activation within the sub-population was designated as the winning BO cell for plotting. Each row in this figure demonstrates maps from a single scale of BO neurons modeled in the hierarchy, with the scales denoted on the right. The arrows of the color wheel in the third row of the plot indicate the figure inside direction in the BO maps. For example, a red pixel signals that the figure lies below that pixel, that is, the pixel is part of that figure’s occluding contour. A close-up of the EBO map for scale 2 responses is shown, with colored arrows indicating figure inside direction. To trace around an object contour in a clockwise direction, follow the directions of the color wheel keeping the direction of arrows to the right of the tracing direction with the slant determined according to the contour pixel color as indicated by the color wheel. Note the disjoint representations in LBO and EBO maps, similar to the luminance and LM maps in Figure 1, but with figure side encodings. The last column shows the joint LBO and EBO maps at each scale, demonstrating how both representation types are needed to give rise to a complete figure ownership representation for this scene.

Comparing mEBO and mLBO maps in each row provides a vivid image of disjoint representations in mEBO and mLBO cells, confirming our previous observations regarding each cell type. As in the joint LM-lum edge map of Figure 1, the joint mLBO-mEBO map in each row of Figure 7 demonstrates the need for both neuron populations to achieve a comprehensive picture of figure ownership in natural scenes; where one neuron population is blind, the other sees ownership direction, providing the signal required for figure-ground organization and to higher visual areas.

## 3 Discussion

We propose the existence of a sub-population of BO cells that respond selectively to equiluminant figure ownership. This proposal, and the predicted response patterns, build on earlier findings of equiluminant neurons in V1 and V4 [26, 5]. We posit that this new kind of border ownership cell is required to both transform equiluminant edge signals and to provide inside-outside information to equiluminant V4 cells with object-centered shape selectivity. Our hypothesis draws on prior work that suggested luminance BO cells as the source of object-side information to V4 neurons [33], by proposing that a parallel mechanism supports the transmission of equiluminant signals.

Color coding documented in 43-64% of BO neurons in earlier studies [59, 15] offers ground for our proposal. Building on these findings, we predict that a sub-population of color-selective BO cells deviates from the commonly observed response pattern in shape-selective neurons, namely, weak responses to low luminance contrast stimuli and stronger activations at high luminance contrast stimuli. Instead, we predict that these neurons exhibit strongest responses at zero luminance contrast with diminishing activation in the absence of color contrast, while maintaining orientation and figure ownership coding similar to that of LBO neuron. With computational modeling, we provided an explanation for how this response pattern emerges in the ventral stream. Our proposed model that was inspired by the RBO network [33], suggests orientation-selective double-opponent neurons [26] as the main source of equiluminance responses in mEBO cells. Early recurrence from mMT neurons in the dorsal stream initiates the divergence of responses to figure on either side of the border. Importantly, dorsal modulation of mEBO cells was based on the observation that the dorsal stream is neither color blind nor color-selective, but color-sensitive [19, 9, 45, 4, 8, 56]. Despite the initial belief on luminance-driven responses in the dorsal stream, several studies noted effects of color in MT responses (see [18] for a review). It is now established that MT neurons indeed respond to equiluminant stimuli, although with reduced strength compared to luminance responses. With color sensitivity in MT neurons, without encoding for the combination of colors, these cells are believed to contribute to computation of object shape [9]. Founded on these results, color-sensitive mMT neurons provide contextual information to our mEBO cells, initiating early figure side encodings followed by further enhancement due to lateral modulations [33, 58].

Our work draws on research that challenged the segregated form and color processing view [24, 21, 31] across different levels (see [17, 18, 49, 48] for reviews). At the input level, Hansen and Gegenfurtner [20] demonstrated that natural scenes contain equiluminant edges that are statistically independent and not rarer than luminance ones (as in the example in Figure 1). At the processing level, findings on equiluminant shape-selective neurons in V1 [26, 16] and V4 [5, 6] suggest that color information is retained within shape-selective neurons across the ventral stream. Additionally, reports on a spectrum of color and form selectivity in ventral areas [16, 25, 27, 46, 7], with a sub-population representing shape solely defined by equiluminant borders, point to an integrated system enabling form perception.

We emphasize that joint color and form coding in EBO neurons does not imply the non-existence of the property binding problem, contrary to the view of Friedman *et al*. [15]. The property binding problem in vision addresses how the visual cortex aggregates independently processed visual features such as shape and color to form a uniform percept (See [57] for a detailed discussion). Friedman *et al*. [15] posed color-selective BO neurons as multiplexing cells, those responding to more than one visual attribute and suggested the joint coding of color and BO features in these neurons resolves the need to solve the binding problem. Although our work builds on their findings, our proposal for EBO aligns with observations of Garg *et al*. [16], *ie*., EBO is a sub-population within a spectrum of selectivities that have BO representation in common. Ultimately, representations across the spectrum, even though not completely independent, might be aggregated and bound in the visual hierarchy to form a unified percept, for example, in figure ground organization. Figure 7 gives one such example. How and where in the visual hierarchy binding of attributes happens is beyond the scope of this work. Here, we remain with our proposal of EBO cells as a sub-population of BO neurons and a hierarchical model explaining how these representations can be achieved in the visual cortex.

The model developed in this work was designed to provide computational evidence for the plausibility of equiluminance responses in BO neurons and to offer a mechanistic explanation of how such representations may emerge in the primate ventral visual stream. To achieve this, we constructed an analytically defined network whose architecture and neuronal properties were directly informed by neurophysiological findings (See Supp. Information for details). Our model includes both luminance and equiluminance subpopulations. Previous computational work has proposed a network in which color-selective neurons represent surface color [32]. Incorporating these representations into the eRBO framework would extend the model into a unified architecture that integrates multiple sub-populations contributing to a coherent object percept in the visual cortex.

Although deep neural networks (DNNs) have been shown to posses certain similarities with primate color representations [42, 13, 1, 52], investigating equiluminant form perception using DNNs remains challenging. Most existing studies rely on DNNs trained on natural RGB images for tasks such as image classification or object segmentation. To reconcile differences between RGB and biologically relevant color spaces, prior work has often involved conversions to color spaces such as DKL for stimulus design or post hoc analysis. However, these transformations can introduce artefacts, particularly when studying equiluminant representations. Converting stimuli that are equiluminant in a biologically plausible color space into RGB inevitably introduces spurious luminance contrasts along object borders, thereby violating the very condition of equiluminance that defines such experiments.

A future direction would be to train DNNs directly on cone activation signals (LMS inputs) to examine whether they can develop color-form representations, including equiluminance ones, analogous to those observed in the primate visual system. In contrast, the present study adopts an analytically specified model based on known principles of color–form processing in the primate brain. This approach enables a controlled and mechanistically interpretable examination of the computational feasibility of EBO cells in the primate ventral stream. With this analytically-defined model, we were able to identify the distinct and complementary response patterns in mLBO and mEBO sub-populations. Our analysis revealed that even though mEBO cells are color-selective [59, 15], their responses extend beyond color-selectivity and point to retaining of equiluminance response properties across the ventral stream. Our model predicts details of border ownership neurons that future experimental research including equiluminant stimuli may test potentially leading to a more comprehensive picture of how object shape is determined in the visual cortex.

## 4 Supplementary Information

### 4.1 Choice of model

The proposed model developed in the present work was intended to provide computational support for the plausibility of equiluminance responses in BO neurons, independent of the mechanisms by which BO signals are generated in the ventral stream. In particular, any model whose neurons encode BO representations could be utilized, conditioned on the plausibility of integrating color stimuli responses in all the neurons contributing to BO responses. Here we chose the RBO network [33] as the underlying model. The RBO network was proposed as a plausible mechanistic model of BO representations in the ventral stream. Specifically, in [33], analysis of the time course of various mechanisms that could potentially provide context to BO neurons led to the suggestion of dorsal modulations as a source of early divergence in BO responses. Experimental results outlined in [33] presented that BO neurons in RBO indeed exhibited similar responses as BO cells in the primate visual cortex [59]. Accordingly, we employed and extended the RBO network to test the hypothesis of EBO representations in the ventral stream.

The RBO network meets the color response criteria. Specifically, in the ventral stream, orientation-selective double-opponent neurons in V1 [26] provide support for modeling color responses in shape processing cells. Similarly, several studies provided evidence on color responses in dorsal pathways [19, 9, 45, 4, 8, 9], suggesting color-sensitivity instead of color-blindness in the dorsal stream [9]. In particular, it was shown that MT neurons respond to equiluminant stimuli [45, 18] and that despite reduced activation strength, their responses were not significantly different from those of ventral areas [18]. Although these neurons do not encode particular hues, magnocellular LGN projections are deemed as the source of color sensitivity in dorsal cells [4]. With evidence supporting color responses in both ventral and dorsal neurons, the RBO network constitutes a plausible computational model for the present work.

### 4.2 Model architecture

In order to test the hypothesis of EBO representations in the ventral stream, we extended the RBO network [33], maintaining its two-stream ventral and dorsal pathway architecture, with signals that meet at border ownership cells. The architecture of the proposed extended model called eRBO is shown in Figure 2. Similar to the RBO network, neurons in eRBO were modeled based on known properties of their biological counterparts in the visual system. Receptive field sizes in the model increased from one layer to the one above [14] with dorsal stream cells having larger receptive fields compared to those of the ventral stream [3]. Additionally, we modeled neurons in both ventral and dorsal streams at four scales, with each scale defining an independent path from that of others. The luminance sub-population in all the model layers were adopted from the RBO network. However, we extended orientation selectivity from horizontal and vertical to 12 orientations in [0*, π*) across both ventral and dorsal pathways. Implementation details and the choice of parameters in luminance neurons can be found in [33].

In eRBO, the overall mechanisms are in common between luminance and color responding cells in both streams. Specifically, in both luminance and color sub-populations in the ventral stream, model simple cells are selective to borders and bars and model complex cells integrate responses of model simple cells [22, 23, 24, 50]. The dorsal pathway models responses of dorsal V1 simple cells and MT neurons to static stimuli [2, 43, 29, 12]. The color-responding sub-population in the dorsal stream is color-sensitive [9]. Therefore, we refer to these neurons as color-sensitive cells. In contrast, ventral neurons responsive to color stimuli were modeled according to equiluminance mechanisms [26], and thus we refer to these cells as equiluminant neurons.

The final layer of eRBO includes model BO neurons that receive feed-forward signals from model complex cells. Responses of mEBO and mLBO neurons in the final layer are multiplicatively modulated by color-sensitive and luminance mMT cells respectively. Dorsal modulation from each neuron’s preferred ownership side results in an initial figure side preference. Then, lateral modulations enhance figure side preferences by iteratively updating mBO responses based on neighboring mBO neuron activations. Our implementation of lateral modulations follows that of the RBO, conducted within individual sub-populations. Unlike in the RBO, we did not perform a scale selection step prior to lateral interactions to aggregate mBO responses across scales. Instead, the four scales of mBO cells are maintained in the final layer and lateral modulations in local neighborhoods are performed independently in each scale.

The following outlines the additional computational components that enable color response modeling in eRBO. For brevity, we omit details common to both eRBO and RBO. We refer the reader to [33] for a full description of the underlying computations and parameter setup.

#### 4.2.1 Cone activations as input

As depicted in Figure 2, input to eRBO are Long (L), Medium (M), and Short (S) cone activations. For any given stimulus presented in RGB, we first converted the input to L, M, and S cone activation using the transformation algorithm proposed by [10]. We adopted and converted the C code provided by the authors to Python. The computed LMS cone activations are then fed to color sub-populations in each stream. Luminance maps that form the input to luminance sub-populations in both ventral and dorsal streams were obtained by linearly combining L and M cone activations as *lum* = *L* + *M*, following [20].

#### 4.2.2 Model luminance sub-populations

The luminance operations in both ventral and dorsal streams begin with a convolution operation on the luminance map to extract orientation features. In the ventral luminance sub-population, selectivity to orientated bar and border patterns at two luminance contrast polarities are modeled. In the dorsal stream, selectivity to orientated bars is modeled in luminance dorsal simple cells whose responses are fed to luminance on- and off-center mMT cells. Details on receptive field sizes, convolution kernels and parameters in luminance cells can be found in [33].

#### 4.2.3 Model equiluminant ventral simple and complex cells

The equiluminant simple cells in the eRBO are modeled according to the findings of Johnson *et al*. [26]. The selectivity profiles of these neurons are demonstrated in Figure 8. As in model luminance simple cells, we modeled selectivity to bar and borders. As shown in Figure 8, both types of selectivity profiles consist of oriented sub-regions with opponent cone contributions within each sub-region. In our model, bar-selectivity in equiluminant simple cells was achieved by convolving input cone maps by a Different of Gaussian (DoG) kernel, followed by a linear combination of resulting maps. Similarly, border-selective cell responses were computed by a weighted combination of cone maps that were convolved with Gabor filters. Specifically, model equiluminant simple cell responses whose receptive field centered at (*x, y*) and was selective to orientation *θ* were obtained by,

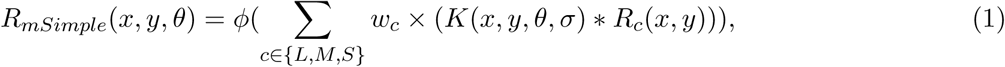

**Figure 8:**
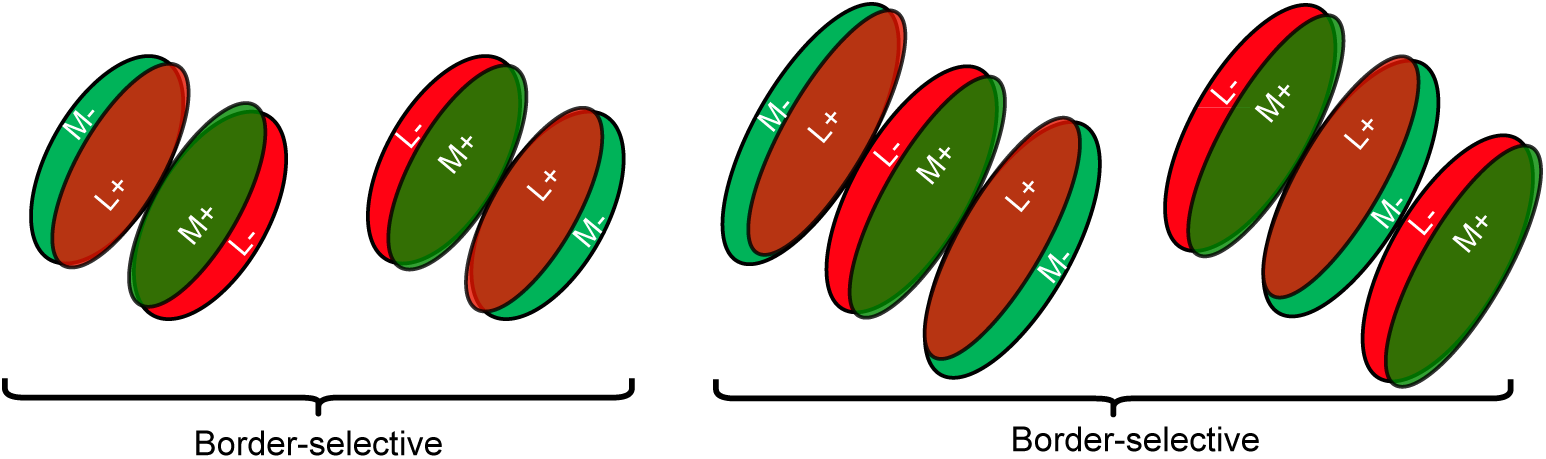
Selectivity profile of modeled equiluminant simple cells in eRBO with bar and border selectivities, following [26] (their Figure 9). Both selectivity types consist of oriented sub-regions with opponent cone contributions. Here, we demonstrated examples with L and M opponent cone contributions. As demonstrated, bar selectivity profiles can be modeled by first convolving cone maps with Gabor filters and then linearly combining them with a negative weight ratio. In a similar manner, border selectivity profiles result from DoG-filtered cone maps that are linearly combined with negative weight ratios.

where *R_c_* represents cone responses with *c* ∈ {*L, M, S*} indicating cone type, ∗ is a convolution operation and *K* is the convolution kernel with parameters *σ*, *w_c_* denotes cone weights and *ϕ* is a linear half-wave rectifier. In this formulation, *K* is either a Gabor or DoG kernel with the same parameters as those of model luminance simple cells. Following [25, 49], we set cone contributions, *w_c_*, to model equiluminant simple cells to have negative ratios as summarized in Table 1. This yields four distinct model equiluminant simple cell types that we named according to their excitatory cone input; for example, L-on for excitatory L input. Responses of model equiluminant complex cells in eRBO are then obtained through a weighted sum of model equiluminant simple cells, with identical weights as those of the model luminance complex cell sub-population.

**Table 1:**
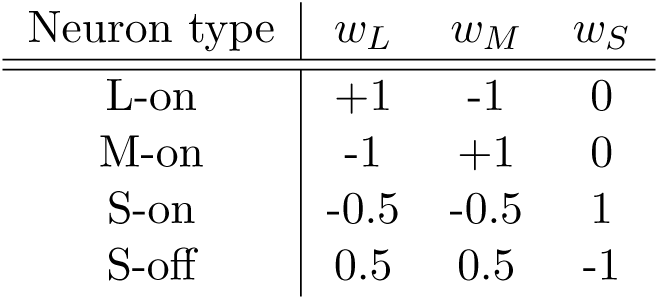
Cone weights to model double-opponent orientation-selective simple cells. Each neuron is identified by the excitatory cone input.

#### 4.2.4 Model color-sensitive dorsal simple V1 and MT cells

Model color-sensitive dorsal simple cells are selective to oriented edges and their responses are obtained by utilizing a DoG kernel in Equation 1, with identical parameters as model luminance dorsal simple cells. The cone weights in these neurons are set to *w_L_* = 0.37*, w_M_* = 0.54*, w_S_* = 0.09. This choice of cone weights follows [8], and aligned with the 1:10 ratio of S to L-M cone response strength in dorsal neurons [4]. Negative cone weights were employed in Equation 1 to model simple cells with inhibitory centers. Finally, color-sensitive mMT neurons, with ON and OFF receptive field profiles, receive their feed-forward input from model color-sensitive dorsal simple cells in the same manner and with identical parameters as those of luminance mMT neurons.

#### 4.2.5 Model equiluminant border ownership cells

In the RBO, dorsal modulation was suggested as the source of initial response divergence to preferred and non-preferred stimuli reported in primate BO cells [59]. In our extended model, mEBO cells receive dorsal modulations from color-sensitive mMT neurons on their preferred side. Then, lateral modulations enhance the difference of responses in mBO neurons with opposite figure side preferences. As in the RBO network, this is achieved by a few iterations (*t* = 10) of relaxation labeling [60, 61], conducted individually within each sub-populations of mBO neurons, *ie*., mLBO-bar, mLBO-border, mEBO-bar, and mEBO-border. Similar to RBO, relaxation labeling among mEBO cells provides local context and deals with uncertainties among neighboring neurons.

In our implementation of relaxation labeling, we utilized the set of dorsally-modulated mBO responses as the initial label for each visual field location (*x, y*). In each relaxation iteration, mBO responses compatible with those of neighboring neurons are strengthened and incompatible ones are weakened. The strengthening or weakening effects are determined based on a compatibility function between labels, here, various mBO selectivities for each visual field location. Formulating a universally applicable compatibility function for the diverse set of mBO selectivities in eRBO was not straightforward. Here, we explain our formulation by first focusing on a simple example.

Imagine a figure border coincides with a visual field location. We consider the set of mEBO cells whose receptive field centers falls on the particular visual field location. Also, we consider these neurons to be border-selective with L and M cone inputs. This particular sub-population of mEBO cells define a set of labels that we categorize based on their polarity and BO direction. Specifically, the border selectivity profiles shown in Figure 8 demonstrate two polarity possibilities for the labels. To determine possibilities regarding BO directions, we define *b* as

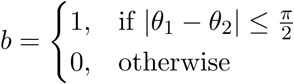

where *θ*_1_*, θ*_2_ denote BO directions for two neighboring neurons and *b*, that we call BO alignment, is a binary variables that signals if the BO directions from the two cells are relatively aligned. With two polarity contrasts and two BO alignment possibilities, four possible label categories emerge and we determine compatibilities within each category.

Figure 9 demonstrates examples of these four possibilities that we consider to define compatibilities. Let *ϕ* = |*θ*_1_ − *θ*_2_| be the angle between the two BO directions. Our goal is to assign high compatibilities to scenarios in which two neighboring cells signal similar representations in a local neighborhood and their BO directions are relatively aligned – same polarity, same BO alignment. All other cases are incompatible as visualized in Figure 9. Here, a variation of cos(*ϕ*) seems an appropriate choice to incorporate in our compatibility function. For example, in the case of same polarity, same BO alignment, cos(*ϕ*) as the compatibility function assigns higher compatibilities to smaller *ϕ*. The cos(*ϕ*) function is also suitable in the opposite polarity, opposite BO alignment case, assigning negative compatibilities according to the angle between BO directions. Similarly, − cos(*ϕ*) ensures incompatibilties determined by the angular difference between BO directions with opposite polarity but aligned BO directions. Finally, same polarity, opposite BO alignment denotes cases in which incompatibilities are achieved by scaling and shifting − cos(*ϕ*) values (−*w* ×cos(*ϕ*)−(1−*w*)). We set *w* = 0.25 in the compatibility function pertaining same polarity, opposite BO alignment to ensure compatibilities between neurons are negative. These four possibilities are summarized in Table 2.

**Figure 9:**
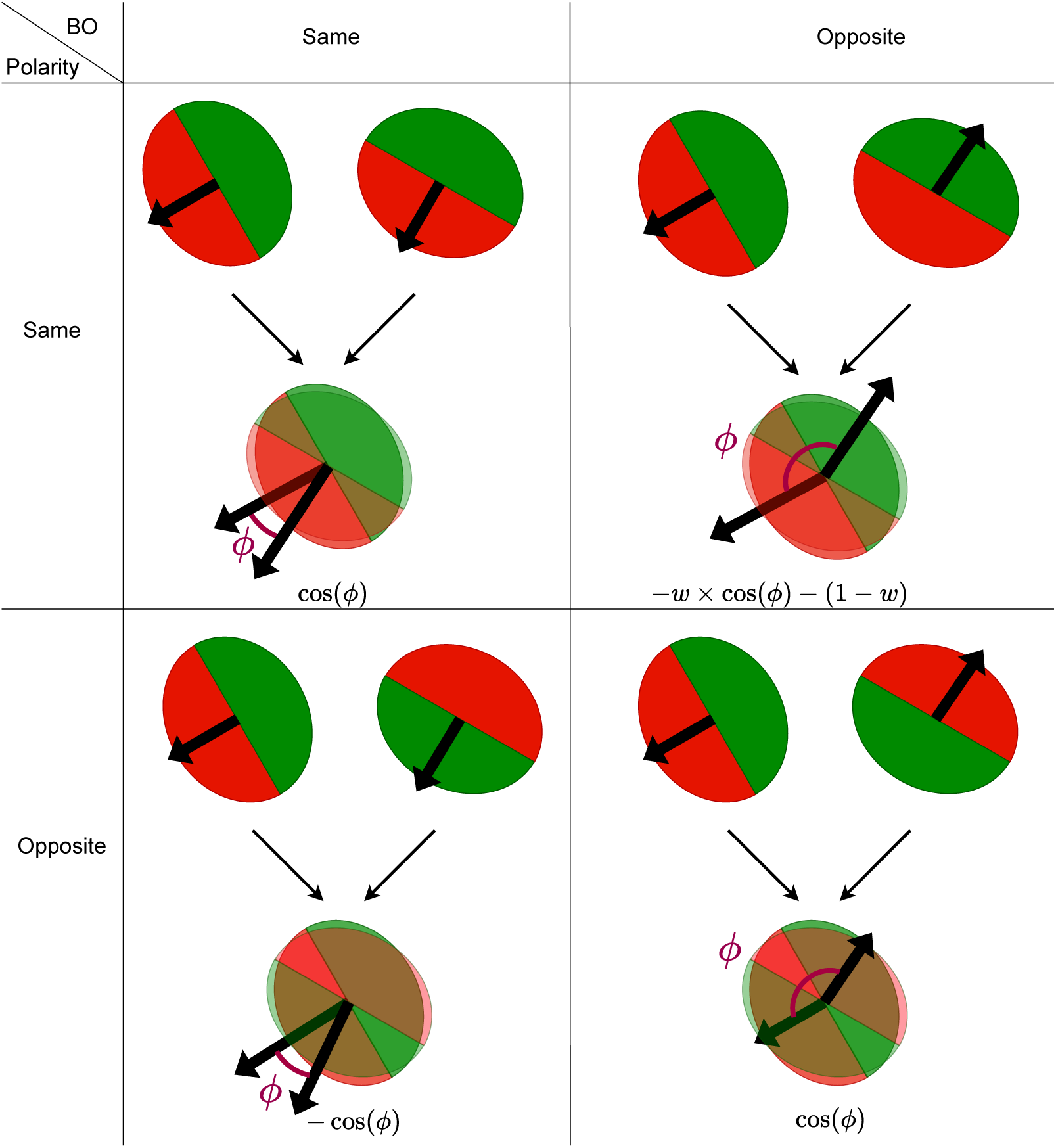
Examples of four possibilities of polarity contrast and BO alignment (*b*) among neurons. The combination of polarity and BO alignment determines the compatibility between the two settings. For example, same polarity and same BO alignment are highly compatible. Therefore, the quantitative value in this case is defined according to the angle between the BO alignment; the smaller the angle, the higher the compatibility. In contrast, when two neurons represent the same polarity contrast but their BO alignments are to opposite directions, they are incompatible. Hence, their quantitative compatibility value is negative.

**Table 2:**
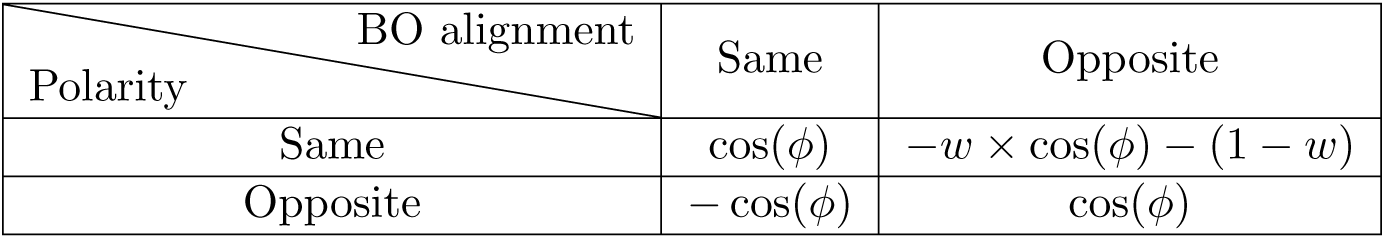
The compatibility function in the relaxation labeling step was determined by first breaking the possibilities into four groups based on polarity and BO alignment among neighboring neurons.

Note that these four possibilities were defined based on binary polarities, *ie*., same or opposite. But in our set of mEBO representations, polarities are not limited to same or opposite - for example, between neurons with L-M and S-LM representations. Therefore, we formulated the compatibility function as a convex combination of four terms defined by the polarity distance, *d*, among the representations,

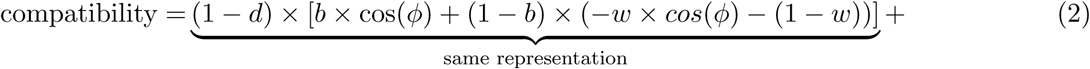

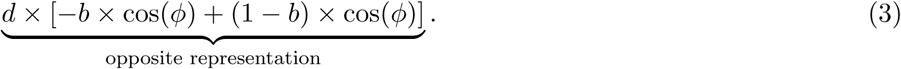

This formulation reduces the compatibility function to one of the cases in Table 3 when *d* = 0 or *d* = 1. For example, *d* = 1*, b* = 0 corresponds to opposite polarity, opposite BO alignment - the bottom right cell in Table 2. However, for *d* ≠ 0, 1, the compatibility is a convex combination of same or opposite representations. In our implementation, we employed polarity distances as defined in Table 3 among mEBO cells. Polarity distances between mLBO neurons were set to 1 or 0 for same or opposite representation respectively, as we employed the same compatibility function for mLBO lateral modulations.

**Table 3:** Polarity distances among four types of mEBO representations utilized in our implementation to determine compatibilities among neighboring cells.

|  | L-on | M-on | S-on | S-off |
| --- | --- | --- | --- | --- |
| L-on | 0 | 1 | 0.5 | 0.5 |
| M-on | 1 | 0 | 0.5 | 0.5 |
| S-on | 0.5 | 0.5 | 0 | 1 |
| S-off | 0.5 | 0.5 | 1 | 0 |

Although we defined the compatibilities and polarity distances intuitively, the compatibility function and the various parameters in determining the compatibilities among labels can be learned for specific domains and applications with availability of relevant data for a more accurate estimate. Finally, in our implementation, updates to a neuron’s label were weighted by the distance of neighboring cells. Local neighborhoods were defined as the set of immediate neighbors in 3 × 3 regions and the distances were determined as,

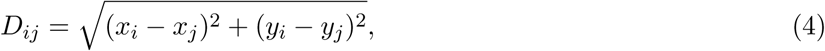

where *D_ij_*denotes the distance between neighbors *i, j*, and (*x_i_, y_i_*), (*x_j_, y_j_*) represent the receptive field centers coordinates of *i, j* neurons.

### 4.3 Additional examples of eRBO on natural images

Figure 10, 11 provide additional examples demonstrating EBO and LBO coding in eRBO. The natural image examples were taken from the McGill Calibrated Colour Image Database [36]. The plots follow the same convention as in Figure 7 in the main article body.

**Figure 10:**
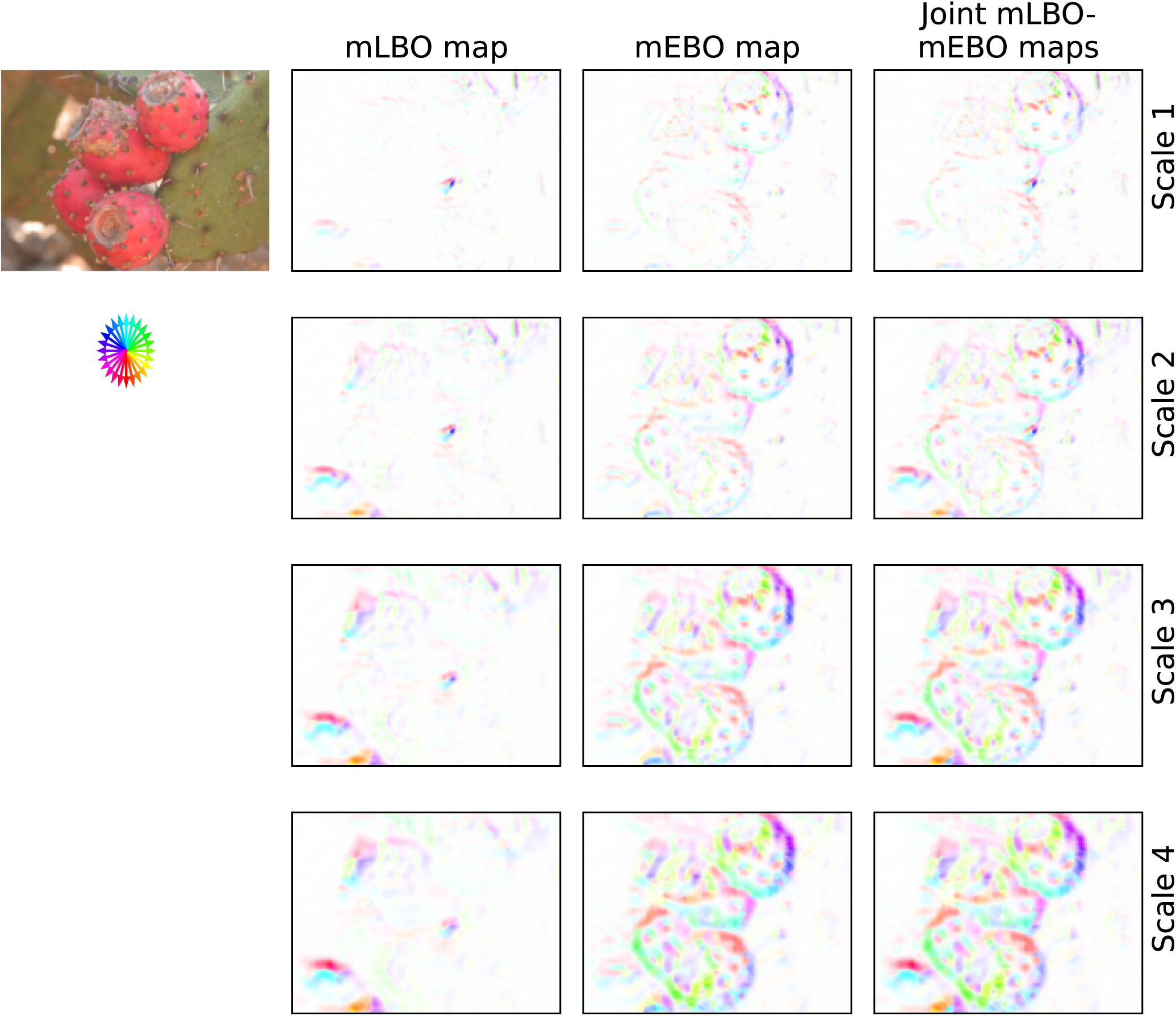
Coding of BO in a natural scene example with the LBO and EBO maps plotted similar to Figure 7 in the main article. Each pixel in these maps represents response of the winning BO neuron whose receptive field centered on the pixel. The BO cell with strongest activation within the sub-population was designated as the winning BO cell for plotting. Each row in this figure demonstrates maps from a single scale of BO neurons modeled in the hierarchy, with the scales denoted on the right. The arrows of the color wheel in the second row of the plot indicate the figure inside direction in the BO maps. To trace around an object contour in a clockwise direction, follow the directions of the color wheel keeping the direction of arrows to the right of the tracing direction.

**Figure 11:**
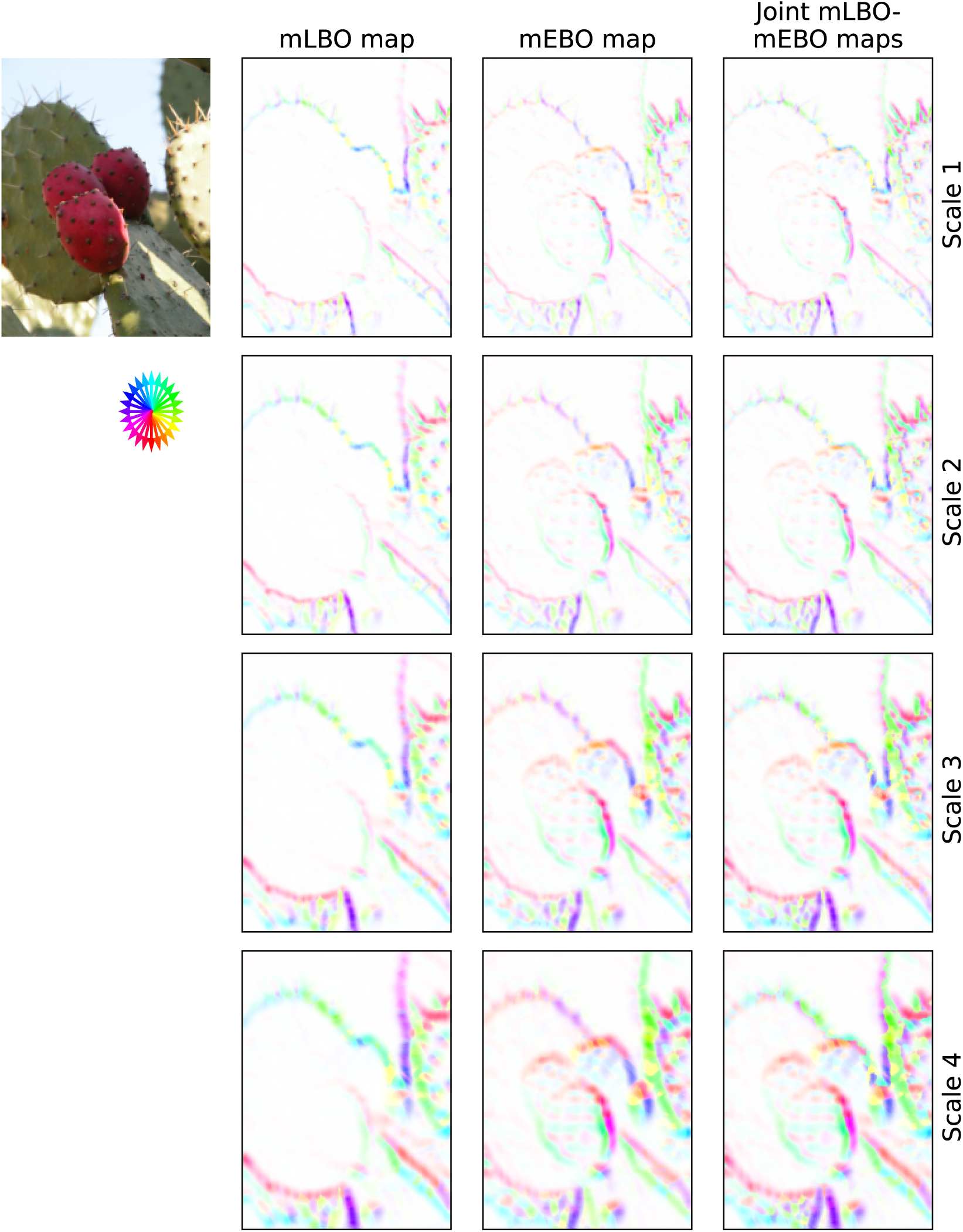
Coding of BO in a natural scene example with the LBO and EBO maps plotted similar to Figure 7 in the main article. Each pixel in these maps represents response of the winning BO neuron whose receptive field centered on the pixel. The BO cell with strongest activation within the sub-population was designated as the winning BO cell for plotting. Each row in this figure demonstrates maps from a single scale of BO neurons modeled in the hierarchy, with the scales denoted on the right. The arrows of the color wheel in the second row of the plot indicate the figure inside direction in the BO maps. To trace around an object contour in a clockwise direction, follow the directions of the color wheel keeping the direction of arrows to the right of the tracing direction.

## Acknowledgments

This work was supported by Air Force Office of Scientific Research Grant FA9550-22-1-0538, Canada Research Chairs Program Grant 950-231659, Natural Sciences and Engineering Research Council of Canada Grant RGPIN-2022-04606.

## Notes

### Competing Interest Statement

The authors have declared no competing interest.

